# Genetic Disruption of System xc- Mediated Glutamate Release from Astrocytes Increases Negative-Outcome Behaviors While Preserving Basic Brain Function in Rat

**DOI:** 10.1101/2022.09.14.504799

**Authors:** Evan M. Hess, Sara Kassel, Gregory Simandl, Nicholas J. Raddatz, Brian Maunze, Matthew M. Hurley, Michael Grzybowski, Jason Klotz, Aron M. Geurts, Qing-song Liu, SuJean Choi, Robert C. Twining, David A. Baker

## Abstract

The impact of CNS disorders is exacerbated by the difficulty in developing safe, effective glutamatergic therapeutics. Synaptic glutamate transmission is vital for neural physiology throughout the brain, which contributes to the vast therapeutic potential and safety risk of glutamatergic therapeutics. Here, we created a genetically modified rat (MSxc) to survey the range of brain functions impacted by the loss of glutamate release from astrocytes involving system xc- (Sxc). Eliminating Sxc activity was not lethal and did not alter growth patterns, activity states, novel object recognition or performance of other simple tasks. In contrast, MSxc rats differed from WT in Pavlovian Conditioned Approach and cocaine self-administration/reinstatement paradigms. Both WT and MSxc rats readily learned that a cue predicted food delivery during Pavlovian Conditioned Approach training. However, WT rats were more likely to approach the food tray (i.e., goal tracking) whereas MSxc rats were more likely to approach the food-predicted cue (i.e., sign tracking) even when this behavior was punished. In the self-administration/reinstatement paradigm, MSxc rats had higher levels of cocaine-primed drug seeking in the absence of altered extinction or cocaine self-administration. These data demonstrate that Sxc-mediated glutamate release from astrocytes regulates non-reinforced and negative-outcome behaviors without altering simple learning or other forms of basic brain function.

## Introduction

Unhealthy behaviors contribute to the severity of addiction, cancer, and other chronic disorders (Verdejo-Garcia et al., 2015; Lange et al., 2017; Harrison and Wefel, 2018; Peters et al., 2019). Although neural circuits influencing behavioral control have been identified (Hyman et al., 2006; Volkow et al., 2017; Luscher et al., 2020), there is a need to identify molecular mechanisms that promote behavioral control and can be safely targeted. This has been challenging, in part, because neuronal targets are typically involved in multiple brain processes, which can impede the development of safe and effective CNS therapeutics. For example, glutamate signaling is often implicated in the pathology and treatment of CNS disorders because it is vital to neural physiology throughout the brain (Vandame et al., 2013; Fairless et al., 2021; Henter et al., 2021), yet few direct, CNS-active glutamatergic therapeutics exist because of the difficulty in obtaining an acceptable safety profile that doesn’t result in excessive reductions in efficacy.

Evolutionary increases in signaling complexity appear to have equipped the brain with mechanisms that have highly restricted or specialized functions in the brain. For example, the expansion of the synaptic proteome resulting from genome duplication enabled more complex regulation of brain function through the introduction of novel paralogues with specialized functions (Emes et al., 2008; Nithianantharajah et al., 2013; Grant, 2016). In support, mutations of *Dlg* paralogues resulting from the first genome duplication (*Dlg* 1 and 4) produce widespread disruptions in CNS function; mutations to *Dlg* paralogues resulting from the most recent genome duplication (*Dlg* 2 and 3) selectively impaired complex cognition without impacting survival rates or simple learning (Nithianantharajah et al., 2013). While the intracellular location of *Dlg* proteins may limit the therapeutic relevance of these findings, leveraging the approach used by Grant and colleagues to use insights from evolution to identify functionally specialized signaling mechanisms could accelerate the development of novel, safe, and effective CNS therapeutics.

A second major increase in signaling complexity resulted from the expansion of the glutamate signaling network. While synaptic glutamate transmission is among the most ancestral forms of inter-neuronal signaling (Ryan and Grant, 2009; Moroz et al., 2014), evolution expanded the molecular basis of glutamate signaling such that it involves a variety of receptors, transporters, and release mechanisms expressed by neurons, astrocytes, and other types of brain cells in vertebrate species (Ryan et al., 2013; Ramos-Vicente et al., 2018; Durkee and Araque, 2019; Moroz et al., 2021), which are expressed by neurons and non-neuronal cells. Hence, a critical question is whether the expanded glutamate signaling network includes functionally restricted or specialized components, as is the case for the synaptic proteome.

Here, we investigated the range of functions impacted by astrocytic glutamate by manipulating system xc- (Sxc) activity. The advantages of Sxc is that it is expressed in astrocytes but not neurons (Ottestad-Hansen et al., 2018), is an evolutionarily-newer glutamate release mechanism (Lewerenz et al., 2013), and the Sxc enhancer N-acetylcysteine has been found to be safe and effective in reducing multiple forms of destructive behavior in humans, including substance use, hair pulling, and self-injury (Grant et al., 2009; Froeliger et al., 2015; Back et al., 2016; Grant et al., 2016; Paydary et al., 2016; Cullen et al., 2018; Squeglia et al., 2018). To begin investigating the range of functions involving Sxc, we created a novel rat model lacking Sxc activity by mutating *Slc7a11*, which encodes the primary functional subunit (xCT) for Sxc. Eliminating Sxc function increased non-reinforced and negative-outcome behaviors in Pavlovian Conditioned Approach and cocaine reinstatement paradigms without altering survival, growth rates, general activity, operant or Pavlovian conditioning, or memory required for object recognition. Hence, altering Sxc activity may impact behavioral control without producing widespread impairments in brain function, which could expedite the development of safe, effective therapeutics for impulse control and related disorders.

## Methods

### Phylogenetic comparisons of Slc7a11/xCT in vertebrates

mRNA protein coding sequences from the *SLC7A11* gene from 115 organisms were retrieved from Genbank (Benson et al., 2009). Partial and isoform sequences were left out of this study. Sequences were aligned by codon using MUSCLE (Edgar, 2004) in the MEGA6 software package (Tamura et al., 2013) and reviewed for accuracy manually. Aligned sequences were run through MEGA6 model selection software using a neighbor joining tree and a maximum likelihood statistical method. Gaps were treated by partial deletion and were not used for computing tree branch lengths. The K2+G (gamma distributed) model was utilized for construction of the phylogenetic tree. Pairwise distances were estimated with uniform rates and partial deletion of gaps. Bootstrap values were generated from 100 replications. A cutoff tree was computed with a minimum bootstrap value of 70. Sequence motifs for specific consensus phosphorylation sites were searched within the alignment and conservational data was manually extracted based on the existence or absence of the site. Determination of sites under positive or negative selection was carried out using the Datamonkey web server (Delport et al., 2010). SLAC, branch-based SLAC, FEL, and an integrative selection analysis (Kosakovsky Pond and Frost, 2005) were carried out using a neighbor joining tree at a 0.05 significance level.

### Animal care and usage

Male Sprague Dawley rats were used in these studies and were obtained from an internal colony of genetically-modified rats generated using a Het/Het breeding approach. Outbreeding involved the use of Sprague-Dawley rats purchased from Envigo, which occurred every three generations. After post-natal day 60, offspring were individually housed. All rats were maintained on a 12 hr/12 hr reverse light/dark cycle with behavioral experiments occurring during the dark cycle. Housing conditions and experimental protocols were approved by Institutional Animal Care and Use Committees at Marquette University and the Medical College of Wisconsin and were carried out according to the US National Institutes of Health guidelines.

### Creation of mutant Sxc (MSxc) genetically-modified rats

Zinc-finger nucleases (ZFNs) were designed targeting the second exon sequence (TGCTAGCTTTTGTTC gagtcTGGGTGGAACTGCTG) to produce small deletions of a limited number of base pairs in the *Slc7a11* gene; capital letters represent binding sites for the individual ZFN monomers, on opposite strands. This mutation was predicted to disrupt Sxc activity because *Slc7a11* encodes the functional subunit of Sxc, xCT. ZFNs were injected into the pronucleus of Sprague Dawley (Crl:SD) rat embryos by pronuclear microinjection of in vitro-transcribed encoding messenger RNAs and the resulting offspring were screened for mutations using a Cel-1 assay and validated by Sanger sequencing as previously described (Geurts et al., 2010) and resulting in single-step, whole-animal disruption of *Slc7a11* (MSxc rats). Deletion of 39 consecutive base pairs (GAGGTCTTTGGTCCCT TGCTAGCTTTTGTTCGAGTCTGG) of exon 2 was confirmed by Sanger sequencing.

### Cell Culture

Astrocyte cultures were generated from WT or MSxc postnatal day 3 rat pups. The striatum was dissected and dissociated using 0.25% trypsin EDTA (Gibco) and cultured in 75 cm^2^ flasks in a humidified incubator at 37°C under 95% O_2_ 5% CO_2_ in Eagles minimum essential medium (Gibco) supplemented with 5% fetal bovine serum/5% horse serum (Atlanta Biologicals), Glutamax (Gibco), and antibiotics/antimycotics (Gibco). To remove debris and non-astrocytic glia, flasks were agitated, and the resulting mono-cell layer was resuspended with 0.25% trypsin EDTA. Cells were counted by hand via a cytometer and seeded in 24-well plates coated with poly-D-lysine and laminin at a density of 200,000 cells per well.

### RT-PCR

Total RNA was extracted from NAc tissue samples using Trizol reagent and was subsequently treated with DNAse (Life Technologies) to remove genomic DNA contamination. RNA purity and quantity were assessed using a Nano Vue Plus spectrophotometer (GE Life Sciences). RNA (1 μg) from each sample was reverse transcribed for PCR (Promega). PCR was conducted using GoTaq DNA polymerase (Promega). Primer sequences were as follows: Slc7a11 (xCT) forward- 5’ AGG GCA TAC TCC AGA ACA CG 3’; Slc7a11 reverse- 5’ TTT AGT CCC ATC AGG TCG TTG 3’; GAPDH forward- 5’ CTC CCA TTC TTC CAC CTT TGA 3’; GAPDH reverse- 5’ ATG TAG GCC ATG AGG TCC AC 3’.

### Glutamate release assay

Striatal astrocytes (DIV14) were incubated for 30 minutes at 37°C in Na^+^-free buffer containing the following: 116 mM choline chloride, 13.4 mM MgSO4, 1.68 mM KH_2_PO_4_, 2.34 mM CaCl_2_, 5.49 mM dextrose, 11.9 mM HEPES, 0.2% choline bicarbonate, titrated to pH: 7.4 with CeOH. This buffer was used to prevent Na^+^-dependent uptake of glutamate. Increasing concentrations of L-cystine 0, 12.5, 25, 50, 100, 200 μM were applied to drive cystine-glutamate exchange by Sxc. Media samples (100 μl) were collected for subsequent glutamate analysis using high performance liquid chromatography (HPLC). Cells were then dissolved in 0.5% SDS and total protein for each well was quantified using the BCA method.

### Glutamate HPLC

The concentration of glutamate was quantified by comparing peak areas from samples and external standards using HPLC coupled to fluorescence detection. A 10 μl sample underwent pre-column derivatization with orthophthaladldehdye (OPA) in the presence of 2-mercaptoethanol using a Shimadzu LC10AD VP autosampler. Chromatographic separation was achieved using a Kinetex XB C-18 (50 × 4.6 mm, 2.6 μm; Phenomenex) and a mobile phase consisting of 100 mM Na_2_HPO_4_, 0.1 mM ethylenediaminetetraacetic acid (EDTA), 10% acetonitrile at a pH of 6.04. Glutamate was detected using a Shimadzu 10RF-AXL fluorescence detector with an excitation and emission wavelength of 320 and 400 nm, respectively. Glutamate content for each sample was normalized to total protein in the respective well and depicted as a net change from baseline.

### Developmental Impact of *Slc7a11* Mutation

To determine the lethality and developmental impact of eliminating Sxc function, we measured mortality rates and body weight gain. Mortality was assessed in two cohorts. First, we monitored survival rates in a large cohort (N>500/genotype/sex) until post-natal day 70, which corresponds to adulthood in rat (McCutcheon and Marinelli, 2009). Next, we monitored survival rates for one year in a smaller cohort (N =8-10/genotype/sex). Body weights were monitored post-weaning until adulthood (i.e., post-natal day 70; N=25-28/genotype).

### Physiology Telemetry

Adult WT and MSxc rats (N=12/genotype) were implanted intraperitoneally with telemetry probes (Mini-Mitter Inc.) to remotely record locomotor activity while in their home cage. Activity data were collected every 5 minutes and averaged in 1-hour bins.

### Novel Object Recognition

During the first two days, adult WT and MSxc rats (N=9/genotype) underwent five-minute habituation sessions, which involved placing the rats in a 50×25cm bedding-free chamber outfitted with a camera. On the third day, rats were placed into the maze for five minutes to become familiarized with two identical objects that were placed in adjacent corners of the chamber. One hour later, rats were placed into the maze for five minutes, which had one familiar object and one novel object. The time spent interacting (looking/sniffing/climbing) with each object was recorded during each session on the third day. The placement and identity of each object were randomized to control for object and spatial preference.

### Pavlovian Conditioned Approach

Adult WT and MSxc rats (N=10 and 9, respectively) underwent autoshaping in operant chambers (Campden Instruments) equipped with touch-sensitive display screens (Horner et al., 2013). One week prior to testing, subjects underwent daily handling and were food deprived to achieve a target weight within 90-95% of their *ad libitum* mass at study onset. In the first of two habituation phases, rats were placed into the chamber until they consumed 20 chocolate-flavored sucrose pellets located in the food tray. This continued once per day until each of the sucrose pellets was consumed within 20 minutes. Next, rats underwent daily 30-minute sessions during which the delivery of a sucrose pellet was paired with the onset of a compound stimulus comprised of a tray light and a 1 second, 3 KHz auditory cue. The pellet and compound cue were delivered on a variable interval (VI) schedule (ranging between 0 and 30 seconds). All rats met the criterion of consuming 40 rewards after one day. Approaches to each side of the food tray were recorded to detect potential side biases, which was defined as ≥ 70% of approaches to one side.

Next, rats underwent Pavlovian Conditioned Approach training over five daily sessions. Each session involved 40 trials (20 CS+ and 20 CS-presentations). The CS+ cue was located on the screen on the rat’s nonpreferred side if a preference was detected during habituation or was randomly determined for rats that did not display a side bias. Each trial was initiated once a 10-40 second VI had passed and the subject broke the infrared beam in the back of the chamber. A visual stimulus (a large white X; Figure 2B) was illuminated on the display screen for 10 seconds. In CS-trials, the intertrial interval began upon termination of the cue. In CS+ trials, the termination of the cue was immediately followed by the compound stimulus (the tray light and a 1-second, 3 KHz auditory cue KHz) and the delivery of one sucrose pellet in the food tray. The intertrial interval (VI, ranging between 10-40 seconds) would then restart. Measures include approaches to each cue screen and the food tray. Physical contacts of the display screens were recorded but occurred too infrequently to be analyzed. After completing 5 daily autoshaping trials, rats underwent 5 sessions of omission testing. These trials were identical to the earlier trials with the sole exception that an approach to the CS+ cue resulted in the omission of the food reward.

### Surgery

WT and MSxc rats that self-administered cocaine were implanted with chronic indwelling catheters. Anesthesia was maintained using 2-2.5% isoflurane using a precision vaporizer during the surgery. A silicon-tubing catheter (0.31 mm ID, 0.64 mm OD) was inserted into the right posterior facial vein until it terminated at the right atrium. The internal aspect of catheter was sutured to the vein. The distal end exited 2 cm posterior to the scapula. The exit port consisted of a back-mounted 22-gauge guide cannula (Plastics One, Roanoke, VA, USA) attached to a polypropylene monofilament surgical mesh (Atrium Medical, Hudson, NH, USA). Rats were allowed to recover for at least 7 days before SA testing during which time they were provided acetaminophen (480 mg/l) in their drinking water. Cefazolin (100 mg/kg, IV) was administered to rats displaying signs of an infection. Post-surgical pain and inflammation was managed using Meloxicam (0.2mg/kg, SC). Catheters were filled daily with a heparin solution (83 i.u./ml) with the distal cannula capped with a closed piece of Tygon tubing whenever the leash/delivery line assembly was disconnected.

### Cocaine Self-Administration and Reinstatement

Adult WT and MSxc rats (N=16 and 14, respectively) were trained to self-administer cocaine (0.5 mg/kg/inf, IV; 2 hr/day) under a fixed-ratio one schedule of reinforcement for 12 daily sessions (Madayag et al., 2010). Rats then underwent extinction training sessions, which were identical to the self-administration sessions except lever presses resulted in saline infusions. Once rats met the extinction criterion of ≤ 15 presses/day, reinstatement testing was conducted. Reinstatement testing involved pretreatment with a low dose of cocaine (3 mg/kg, IP) ten minutes prior to testing and recording lever pressing for 2 hours. This dose was selected because it is typically ineffective in producing reinstatement in WT rats, which thereby enabled us to assess increase reinstatement susceptibility in MSxc rats.

### Experimental Design and Statistical Analysis

Investigators were blinded to genotype for all behavioral procedures. All data are presented as means ± standard error of the mean with individual data points for independent samples and paired lines for dependent samples overlaid. Statistical analyses were conducted using SPSS (v 14.0). Statistical significance was determined using a mixed model, repeated-measures ANOVAs varying Genotype (MSxc vs WT) as the between-subject factor and session (1-5) as the within-subject factor. Fisher’s least significant difference tests or Student’s t-test were used to make planned pairwise comparisons when there were only two means to compare or when ANOVAs revealed an appropriate significant main effect or interaction. The Holm-Sidak procedure was used to contain family wise α level at 0.05 for multiple *post hoc* comparisons. Effect sizes were calculated using partial eta-squared (η_p_^2^; ANOVA main effects and interactions) and Cohen’s d for important pairwise comparisons (Student t-tests and Fisher LSD comparisons). Unique analytical details for each experiment, such as number of rats, degrees of freedom, and follow-up statistical analysis are reported in the corresponding results section.

## Results

*Slc7a11*/xCT arose following the divergence of protostomes and deuterostomes (Lewerenz et al., 2013), which establishes it as an evolutionarily new component of glutamate signaling. To better understand the evolution of Sxc, we conducted a phylogenetic comparison of xCT across 115 vertebrate species which were distributed across the major classes of vertebrates (i.e., Chondrichthyes or cartilaginous fish, Osteichthyes or bony fish, Reptilia, Aves, and Mammalia). We found that *Slc7a11*/xCT is expressed by every vertebrate species examined.

A single likelihood ancestor counting (SLAC) analyses of *Slc7a11*/xCT revealed that it has been under negative selective pressure. The majority of codons (391/569) displayed significantly more nucleotide mutations that did not alter amino acid sequence (synonymous mutations) than mutations that altered amino acid sequence (nonsynonymous mutations; dN/dS ratio p<0.05; figure 1b). The genetic divergence of xCT resulted in a gene tree (figure 1c) that broadly matched phylogenetic relationships among sampled vertebrate species (Irisarri et al., 2017). These findings suggest that while Sxc is a relatively newer release mechanism, it likely encodes a type of signaling that is functionally beneficial to vertebrates.

**Fig.1.**
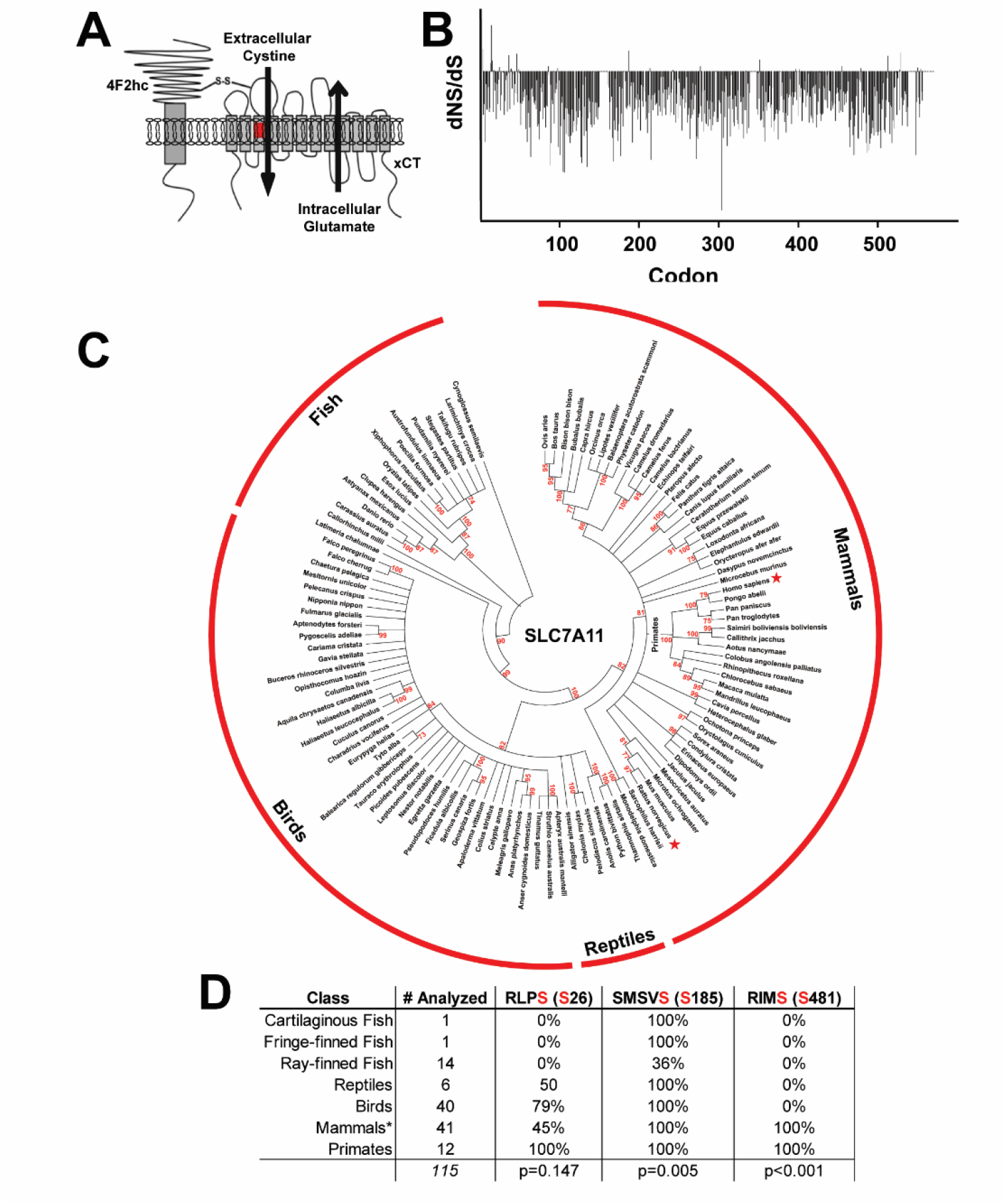
*Slc7a11* is expressed in every vertebrate species examined and is under significant negative pressure. (A). The schematic depicts the two proteins that comprise Sxc, 4F2HC and xCT, and that Sxc releases glutamate through the heteroexchange of extracellular cystine. (B) A SLAC plot illustrating the difference in the number of nonsynonymous (dNS) and synonymous (dS) mutations at each of the Slc7a11 codons across 115 vertebrate species. Most codons (391/569) had significantly more synonymous mutations (i.e., nucleotide mutations that did not alter amino acid sequence) than nonsynonymous mutations, which indicates that the Slc7a11/xCT has been under negative selection pressure. (C) A phylogenetic tree illustrating the xCT protein coding sequence relationship across 115 vertebrate species. (D) The percent of species within vertebrate classes that express designated xCT residues. The p values reflect the outcome of SLAC analyses, which determines whether the site was under selective pressure.

Next, we eliminated Sxc function by creating a novel genetically modified rat model. Zinc-finger nuclease technology was used to excise 39 consecutive base pairs from exon 2 of the gene encoding xCT, *SLC7A11* (figure 2A). The result of this mutation was the complete elimination of cystine-evoked glutamate release in cultured striatal glia obtained from MSxc rats (figure 2B). An ANOVA performed on glutamate release revealed an interaction between genotype and cystine concentration (F_5,79_=2.67, p=0.028, η_p_^2^ = 0.145). Analyses of cystine-evoked glutamate release in WT produced a main effect of cystine (F_5,63_=7.58, p<0.0001, η_p_^2^ = 0.376), with significant increases in glutamate occurring at every nonzero concentration of cystine (Holm-Sidak, the p value for the first comparison = 0.029, the p values for all other comparisons were < 0.0003; all Cohen’s D values > 1.53). In contrast, a significant increase in glutamate was not observed in MSxc-generated cells at any concentration of cystine (F_5,16_=1.39, p=0.280, η_p_^2^ = 0.303). Consistent with these data, we also found that NAc tissue from adult MSxc rats were devoid of xCT (*slc7a11*) mRNA (Fig. 2C). Collectively, these data demonstrate that Sxc function is absent in MSxc rats, which renders this model an excellent tool to explore the functional role of Sxc in behavioral control.

**Figure 2.**
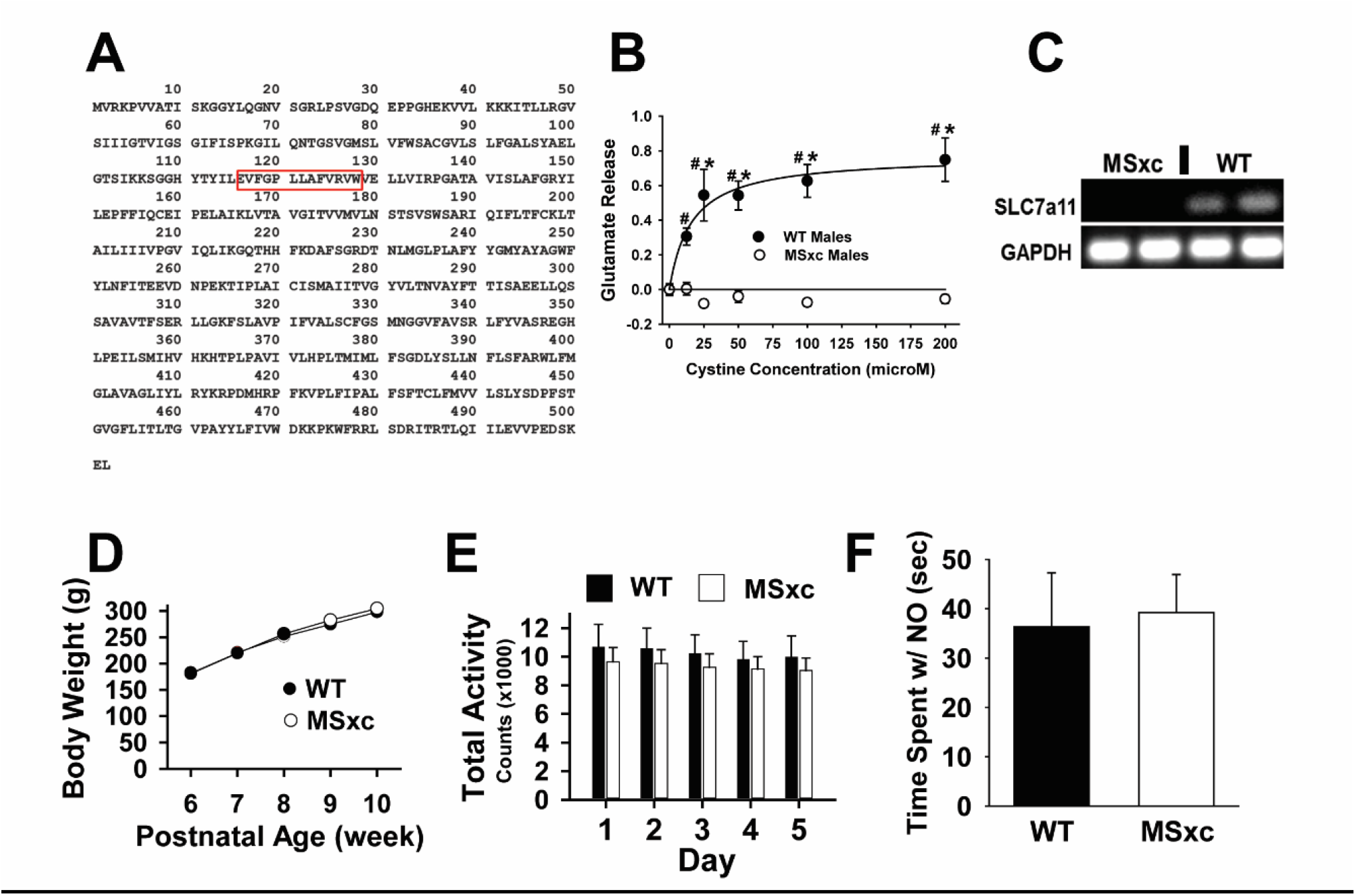
Mutating xCT protein eliminates Sxc function but does not produce generalized impairments in brain function. (A) The 13 amino acids deleted from rat *Slc7a11* are indicated by the red box. (B) Cystine application evokes increase in extracellular glutamate (mean ± SEM) in cultured striatal glia generated from WT but not MSxc rat, indicating a functional knockout of Sxc in MSxc tissue. * Specifies a significant difference from WT rats at the corresponding cystine concentration (Holm-Sidak, p<0.05). # Indicates a significant difference from glutamate content at 0 cystine within genotype. (C) xCT mRNA was detected in NAc tissue punches obtained from WT but not MSxc rats. (D-F) A loss of functional Sxc activity in MSxc rats did not result in altered growth rates (D), home cage activity measured using telemetry probes (E), or time spent with a novel object in the object recognition task (F).

### Basic brain function is unaltered in rats with genetic disruption of Sxc

To investigate the physiological and behavioral impact of loss of Sxc function we first examined genotypic differences in developmental mortality rates, post-natal increases in body weight, exploratory behavior, and simple associative learning which could indicate the range of brain function reliant on Sxc. We did not observe any fatalities in either WT or MSxc rats up to week 10 in a large cohort (N>500/genotype/sex) or during a period of one year in a second (N=8-10/genotype/sex). Growth rate to adulthood and home-cage locomotor activity also did not differ between genotypes. Analysis revealed no genotypic effects on growth rate (figure 2D; n=25-28/genotype; effect of genotype F_1,51_=0.33, p>0.05; week: F_4,204_=22.18, p<0.001; genotype x time: F_4,204_=4.20, p<0.001; p> 0.05 when comparing genotype at each week), or home-cage locomotor activity (figure 2E; n=12/genotype; genotype x day F_7,91_=0.458, p>0.05; genotype F_1,13_=0.331, p>0.05; day F_7,91_=2.759, p<0.05).

Additionally, there were no genotypic differences in time spent exploring a novel object (figure 2F; n=9/genotype; t_16_=0.216, p>.05). These results demonstrate that MSxc rats do not exhibit maladaptive traits in physiology nor deficits in performing a basic memory task. Therefore, subsequent behavioral deficits are not likely to be caused by disruption of these more basic learning or physiological processes.

### Associative learning is unaltered in rats with genetic disruption of Sxc

Next, we used a modified Pavlovian Conditioning Approach task to investigate the impact of diminished Sxc activity on behavioral control. In this paradigm, subjects often display a dominant pattern of approach behavior directed toward a reward-predictive stimulus (i.e., sign tracking) or towards the food tray where the reward will soon be delivered (i.e., goal tracking). When goal tracking predominates, it is thought to reflect dominant behavioral control from executive functions such as response or attentional control; whereas, when sign tracking predominates, it is thought to reflect behavior that is controlled by lower-order processes such as incentive salience or reward (Lovic et al., 2011; Koshy Cherian et al., 2017; Kuhn et al., 2018; Sarter and Phillips, 2018). Importantly, our procedure was modified to minimize neural processes contributing to CS+ approach behavior and the apparent magnitude of sign tracking (Moore, 2004), we used an illuminated CS+ rather than the insertion of a lever which can trigger predatory or consummatory behaviors directed at the lever independent of the learned incentive properties.

Male WT (n=10) and MSxc (n=9) rats were first trained to associate the presentation of a visual stimulus (e.g., CS+) with the impending delivery of food into the food tray. In addition, rats were also trained that a second stimulus functioned as a CS-since it did not predict the delivery of food into the food tray. The ability of WT and MSxc rats to form this association can be determined by analyzing head entries into the food tray during presentation of each stimulus (figure 3). An ANOVA conducted on percent of trials with a tray entry during CS+ presentation revealed a main effect of session (F_4,68_=24.019, p<0.001, η_p_^2^ = 0.586) with no main effect (F_1,17_=0.022, p=0.883) nor an interaction (F_4,68_=0.268, p=0.898) involving genotype. Subsequent post hoc analyses revealed the differences in tray entry between session 1 and sessions 2-5 to be significant (figure 3D; independent of genotype, Holm-Sidak, p values <0.002; Cohen’s D values >0.925). In contrast, there were no significant changes in the percent of trials with a CS-tray entry across training sessions (Figure 3E; session, F_4,68_=1.018, p=0.405; genotype, F_1,17_=1.205, p=0.288; session x genotype, F_4,68_=0.345, p=0.847). These data further establish that loss of Sxc does not alter a rat’s capacity to form and discriminate simple Pavlovian associations. Together, these results indicate that both WT and MSxc rats assign incentive value specifically to the CS+ and subsequently approach the food tray at similar rates.

**Figure 3.**
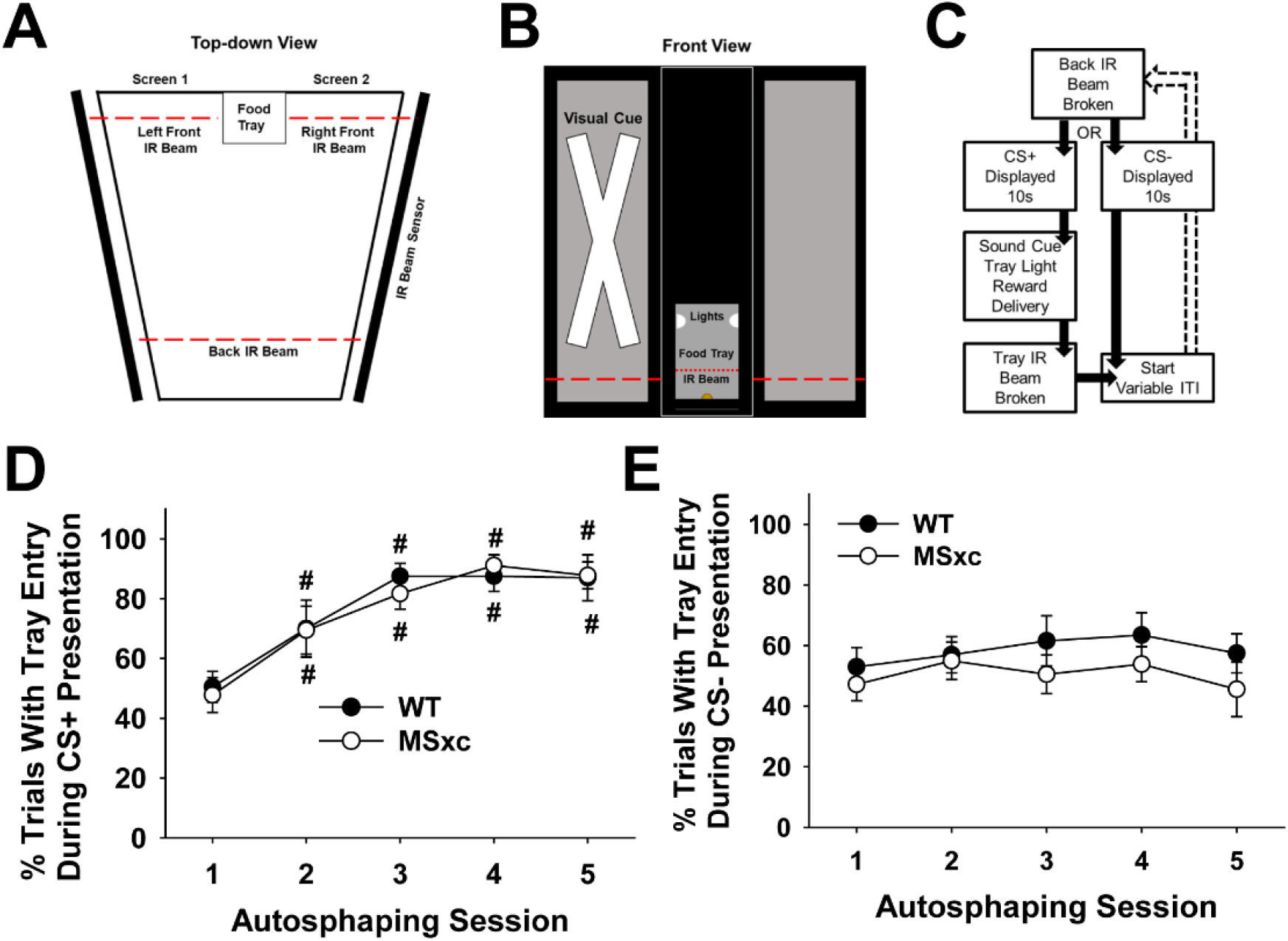
Pavlovian conditioning is not disrupted in MSxc rats. (A-B) An illustration of the Pavlovian conditioning chamber from a top-down and back-front perspective. Red lines depict the location of infrared beams. The large x represents the visual stimulus used as either the CS+ or the CS-. (C) The sequence of events occurring in each trial. (D-E) Figures depict the percent of trials (mean ± SEM) with a tray entry during the presentation of CS+ (D) or CS- (E). # Indicates a significant difference from session 1 (Holm-Sidak, p<0.05).

### Sign-Tracking is augmented in rats with genetic disruption of Sxc

Given that WT and MSxc rats are equally adept at using a predictive cue to approach a goal, we next examined the percent of trials that subjects approached the visual CS+ (figure 4). Although both MSxc rats and WT rats approached the CS+ similarly during session 1, repeated training elicited significantly more CS+ approach by the MSxc but not by WT rats. This observation is supported by a significant interaction between genotype and session (F_4,68_=4.341, p=0.003, η_p_^2^ = 0.203). An analysis across sessions of the percent of trials with a CS+ approach revealed that WT rats did not change CS+ approach behavior across the five sessions (F_4,32_=0.439, p=0.779). In contrast, MSxc rats had significantly more trials with a CS+ approach across sessions 3 and 4 (figure 4A; F_4,32_=6.57, p=0.001; η_p_^2^ = 0.451) compared with session 1 (Holm-Sidak, p values ≤0.014; Cohen’s D values > 1.650; note, session 5, Cohen’s d= 0.89; p=0.034 did not meet the p value adjusted by the Holm-Sidak test). Furthermore, MSxc rats also reliably approached the CS+ more frequently per session (figure 4B). ANOVA revealed a significant two-way interaction between genotype and session F_4,68_=6.571, p=0.0001, η_p_^2^ = 0.279). WT rats did not change the frequency of their CS+ approach behavior across sessions (F_4,36_=0.593, p=0.670), but MSxc rats increased the total number of CS+ approaches across sessions (F_4,32_=5.893, p=0.001, η_p_^2^ = 0.424; Holm-Sidak, sessions 3 and 4 p<0.019, and session 5 p=0.021; Cohen’s D=1.246; figure 4B). Pairwise post hoc t-tests across genotype confirmed that while there were no differences during session 1 (t_17_=0.62, p=0.5471), MSxc rats committed significantly more CS+ approaches compared with WT rats on the last training day (t_17_=2.396, p=0.028, Cohen’s D=1.085). Together, these results suggest that the loss of Sxc predisposes rats to approach reward predictive cues.

**Figure 4.**
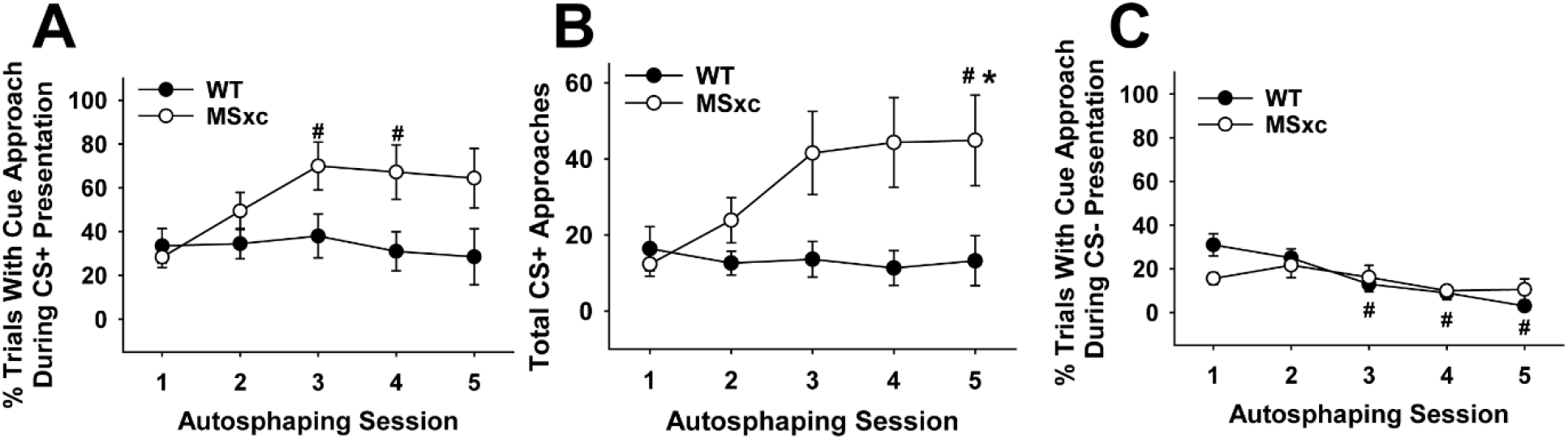
MSxc but not WT rats increase CS+ approach behavior following repeated testing. (A) Data depict the percent of trials (mean ± SEM) with an approach to the CS+ (B) total CS+ approaches and (C) percent of trials with an approach to the CS- across the five training sessions. # Indicates an effect of session relative to session 1 (Holm-Sidak, p<0.05). * Indicates a difference from WT within the indicated session (Holm-Sidak, p<0.05).

### Cue discrimination is intact in rats with genetic disruption of Sxc

In Pavlovian Conditioned Approach designs, rats learn to approach the CS+ but not the unrewarded CS-. Here, WT rats reduced CS- approach across sessions as expected. Interestingly, MSxc rats did not decrease CS- approach to the same extent, instead maintaining stable low-levels of CS- approach (figure 4C). However, both groups showed intact cue discrimination learning. In contrast to their stable CS+ approach behavior, WT rats learned to further suppress CS- approach across training sessions. MSxc rats, on the other hand, increased CS+ approach behavior while maintaining stable and low levels of CS- approach across training (figure 4C).

In support, ANOVA on the percent of trials with a CS- approach confirmed this observation with a significant 2-way interaction between genotype and session (F_4,68_=2.947, p=0.026, η_p_^2^ = 0.148). Follow-up ANOVAs conducted on the groups separately revealed a significant effect of session for the WT (F_4,36_=11.225, p<0.0001, η_p_^2^ = 0.555), but not MSxc (F_4,32_ =1.579, p=0.204, η_p_^2^ = 0.165) rats. Post hoc tests indicated that the percent of trials with a CS- approach was significantly reduced during sessions 3-5 for the WT rats (Holm-Sidak, p values <0.007; Cohen’s D values > 1.642) but remained unchanged in the MSxc rats. Therefore, unlike WT rats, MSxc rats increased approach to the CS+ (figures 4A-B) and maintained a stable low level of approach to CS- (figure 4C).

### MSxc rats display punished cue approach behavior

Increased CS+ approach behavior, sign-tracking, exhibited by the MSxc rats could be considered maladaptive if it were to occur at the expense of reward consumption. However, MSxc rats, as observed in figure 3, approached the goal tray at the same rate as WT controls. This ability to use environmental cues to predict biologically salient events and approach a primary reward is adaptive. Nevertheless, the MSxc rats approached the CS+ significantly more than the WT rats. This pattern of excess CS approach is not required to obtain the reward and can lead to negative outcomes. However, sign-tracking observed in Pavlovian Conditioned Approach can be made to be unambiguously maladaptive by using a punishment design. Hence, we modified the procedure to include punishment of CS+ approach by omitting the food reward on trials with a CS+ approach.

Indeed, CS+ approach behavior persisted in MSxc rats under punishment conditions (figure 5A-B) while goal approach was similar. This observation was supported by ANOVAs revealing no significant changes in the percent of trials with a goal-tray entry elicited by the CS+ (omission session, F_4,68_=1.074, p=0.376; genotype, F_1,17_=1.427, p=0.249; session x genotype, F_4,68_=0.506, p=0.732) or CS- (omission session, F_4,68_=1.213, p=0.314; genotype, F_1,17_=0.905, p=0.355; session x genotype, F_4,68_=0.736, p=0.571). CS+ approach behavior, on the other hand, remained significantly elevated in MSxc rats despite punishment. Although the percent of the trials with a CS+ approach was elevated marginally by disruption of Sxc (figure 5A), total CS+ approaches across omission sessions were significantly increased (figure 5B). In support, an ANOVA conducted on the percent of trials with a CS+ approach yielded only a significant main effect of session (session: F_4,68_=9.076, p<0.001, η_p_^2^ = 0.348; genotype, F_1,17_=2.705, p=0.118; session x genotype, F_4,68_=0.663, p=0.620). Post hoc tests indicated that the percent of trials with a CS+ approach significantly decreased across omission sessions 2-5 for both groups overall (Holm Sidak, p<0.002; Cohen’s D>0.393). Nevertheless, an ANOVA conducted on the total number of CS+ approaches during omission sessions revealed a significant main effect of genotype (F_1,17_=6.187, p=0.024, η_p_^2^ = 0.317) and session (F_4,68_ =7.881, p=0.012, η_p_^2^= 0.317) indicating that MSxc rats maintained a significantly higher rate of CS+ approach behavior compared with WT rats even while significantly reducing CS+ approach behaviors across the omission sessions. This observation was supported by post hoc tests of the main effect of omission session that show total CS+ approaches were reliably reduced during omission sessions 2-5 compared with session 1 (figure 5B: Holm Sidak, p<0.01; Cohen’s D>0.292).

**Figure 5.**
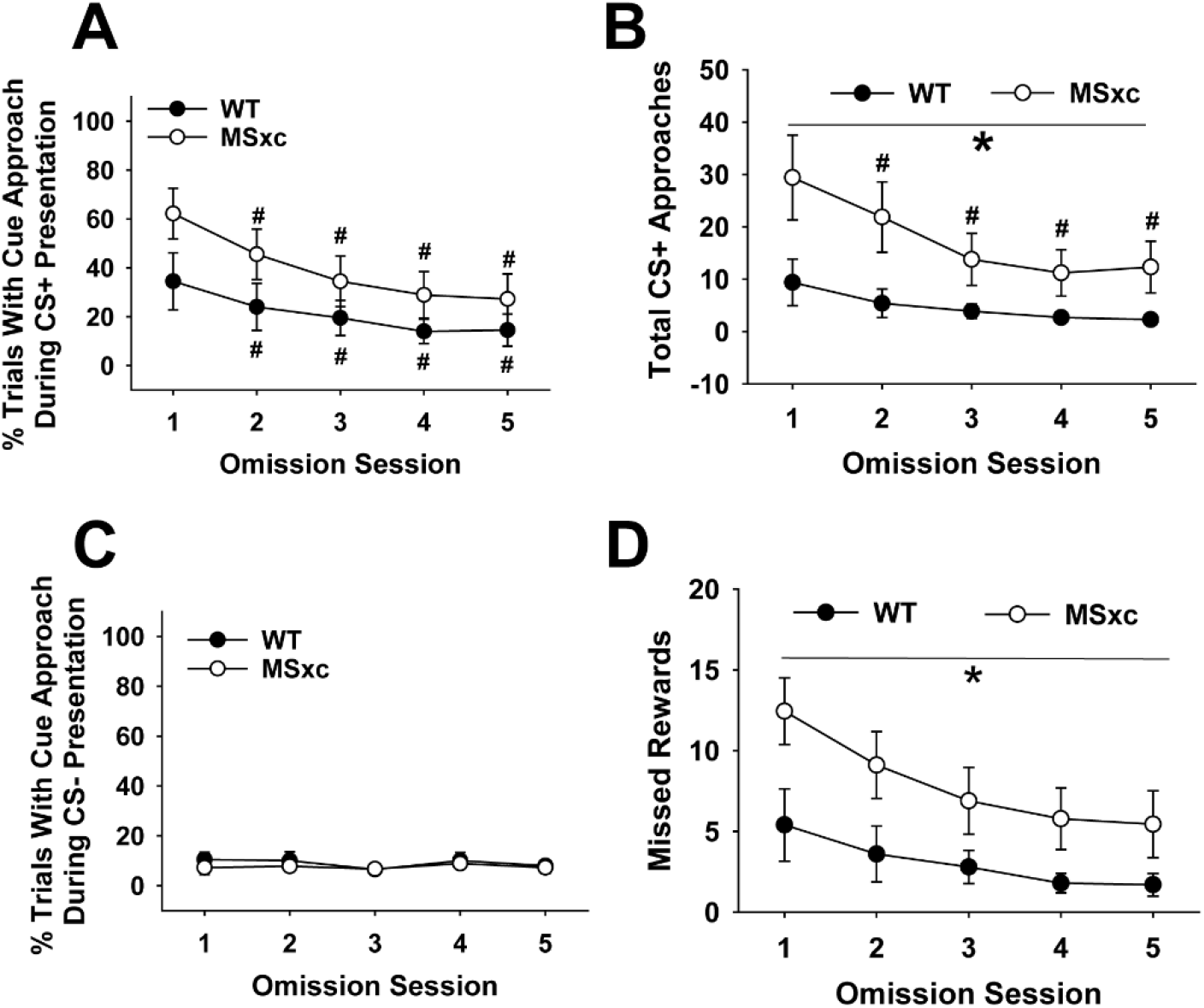
Persistent CS+ approach behavior in MSxc rats persist despite the punishment of this behavior during omission training. (A) Data depict the percent of trials (mean ± SEM) with a CS+ approach, (B) total CS+ approaches per session or (C) the percent of trials with a CS- approach during omission training. (D) These data reflect the total number of missed rewards during approaches toward CS+ during omission training. # Indicates an effect of session relative to session 1 (Holm-Sidak, p<0.05). * Indicates a significant difference from WT (ANOVA, p<0.05).

Critically, this resistance to punishment exhibited by the MSxc rats while sign-tracking lead to a significant failure to obtain the food rewards across the omission sessions (figure 5D). An ANOVA conducted on the total number of missed food rewards revealed significant main effects of genotype (F_1, 17_=5.86, p=0.027, η_p_^2^=0.26) and session (F_4, 68_=8.317, p=0.00002, η_p_^2^=0.329) but no interaction (F<1). Post hoc tests of the session effect indicate a reliable decrease in the number of missed rewards across sessions for both groups (Session 1>Sessions 2-5). Despite this shared decrease, the MSxc rats missed significantly more food rewards under punishment conditions compared with WT rats (Average total missed rewards ± SEM: MSxc= 39.67 ± 8.56 vs WT= 15.3 ± 5.67; Cohen’s d= 1.10). Collectively, these data indicate that punishment of CS+ approach behavior by reward omission does not reduce the increased sign-tracking behavior exhibited by the MSxc rats. Therefore, failure to suppress the perseverative CS+ approach behavior then directly leads to a negative outcome, the loss of food.

In contrast to the increased CS+ approach behaviors, reward omission had no discernable effect on approach to the CS- (figure 5C). In support, there were no significant main effects or interactions when analyzing percent trials with a CS- approach (Figure 5C; session, F_4,68_=0.583, p=0.676; genotype, F_1,17_=0.348, p=0.568; session x genotype, F_4,68_=0.192, p=0.942) or the total number of CS- approaches during omission training (Data not shown; session, F_4,68_=0.870, p=0.487; genotype, F_1,17_=0.523, p=0.479; session x genotype, F_4,68_=0.723, p=0.579). Taken together, these findings highlight the selective nature of the deficit imposed by genetic disruption of astrocytic glutamate release. Impaired Sxc promotes negative outcome behaviors such as increased sign-tracking and resistance to punishment.

### Genetic disruption of Sxc increases drug induced reinstatement of drug seeking

The increased CS+ approach behaviors exhibited by MSxc rats during omission training could indicate that a loss of Sxc may increase the risk for other maladaptive behaviors that are characterized by compulsive or perseverative behavior like drug seeking (Tunstall and Kearns, 2015; Versaggi et al., 2016). If disruption of Sxc leads to a general disruption of top-down control of behavior and/or enhanced incentive sensitization, then MSxc rats would be predisposed to developing compulsive drug seeking. To determine the extent to which Sxc disruption affects drug seeking behaviors, WT and MSxc rats were trained to self-administer cocaine. Drug seeking was then extinguished and evaluated during a reinstatement paradigm. Loss of Sxc had no effect on the acquisition or maintenance of cocaine self-administration. First, WT and MSxc acquired cocaine self-administration at the same rate across days (figure 6a, t_28_ =-1.046, p=0.305) revealing again that these rats do not differ on simple forms of learning. Next, the number of infusions self-administered by WT and MSxc rats during daily 2-hr sessions were comparable (figure 6B). Analyses of this data revealed a main effect of session (F_11,308_=2.237, p=0.013, η_p_^2^ = 0.074) in the absence of a significant main effect (F_1,28_=0.002, p=0.962) or interaction involving genotype (F_11,308_=1.062, p=0.392). Finally, there were no significant differences in intake relative to session 1 that would indicate an escalation of drug intake in either WT or MSxc rats (Holm-Sidak p values >0.106). These data indicate that a loss of Sxc does not alter sensitivity to the reinforcing properties of cocaine in MSxc rats.

**Figure 6.**
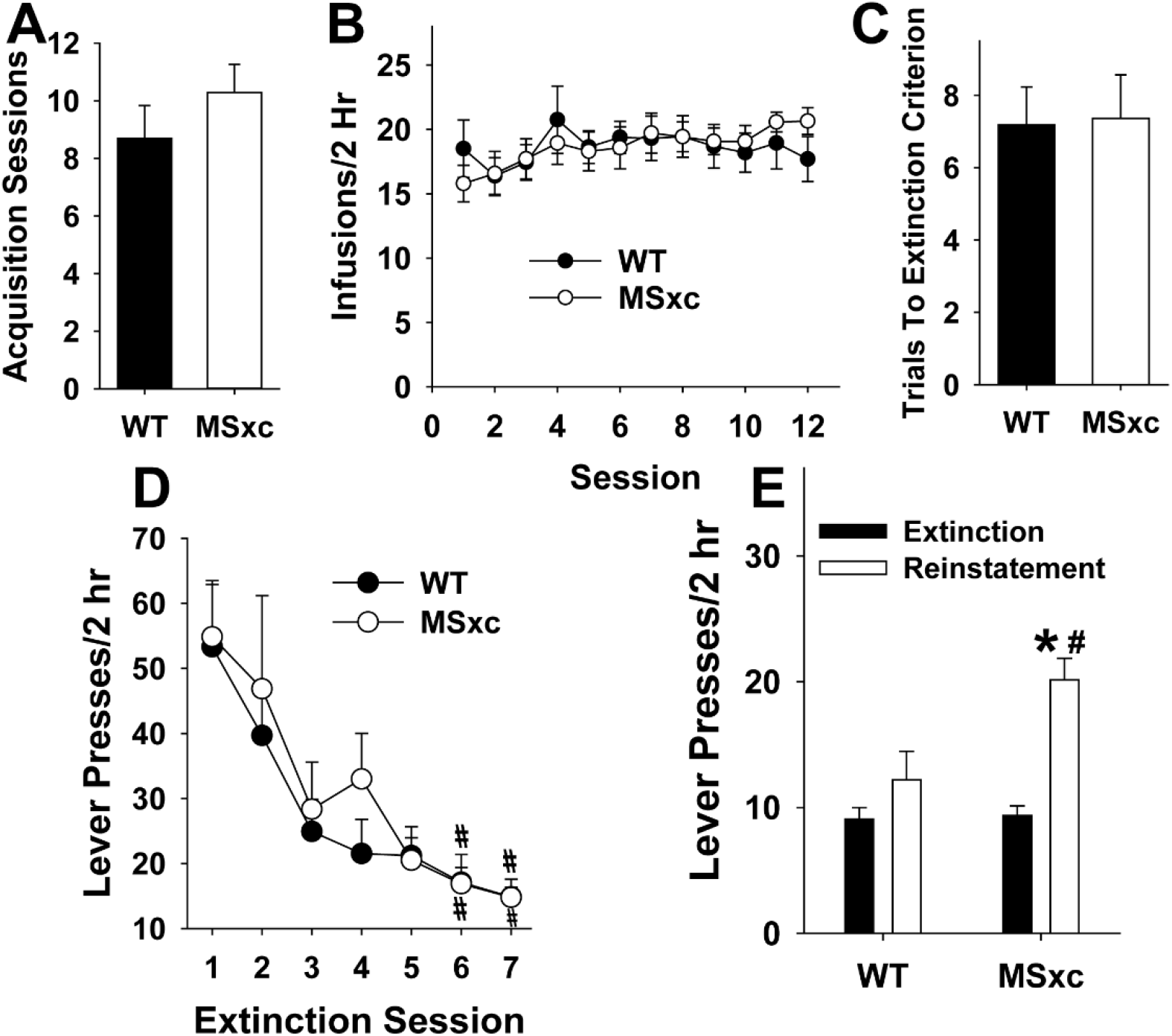
MSxc rats acquire, maintain, and extinguish cocaine self-administration normally but are more vulnerable to cocaine-primed reinstatement. (A) Data depicts the number of training sessions (mean ± SEM) needed to reach the acquisition criterion for cocaine self-administration (0.5mg/kg/infusion (B) Here, we depict the number of cocaine infusions received during the 12 maintenance sessions. (C) These data display the number of extinction training sessions needed to meet the extinction criterion (i.e., fewer than 15 active lever presses in a 2-hour session for two sessions). (D) Here, we depict the pattern of reduced responding between WT and MSxc across the first seven extinction sessions. (E) This figure depicts the number of active lever presses during the last day of extinction and during the cocaine reinstatement test (3 mg/kg, IP). # Indicates an effect of session relative to session 1 of the designated experimental phase (Figure 6D) or the last extinction session within genotype (Figure 6E; Holm-Sidak, p<0.05). * Indicates a genotypic difference during reinstatement (Holm-Sidak, p<0.05).

Likewise, there was no effect of genotype on extinction learning (figure 6C-D). Analysis of this data did not reveal an effect of genotype in the number of trials needed to achieve extinction criterion (Figure 6C; t_28_=-0.107, p=0.915) or in the pattern of reduced responding across the first seven extinction sessions (figure 6D; genotype, F_1,10_=0.100, p=0.758). A main effect of extinction session was present (F_6,60_=7.624, p<0.001, η_p_^2^ = 0.433), but there was no interaction between session and genotype (F_6,60_=1.227, p=0.305). Post hoc analyses revealed that the number of operant responses on the previously cocaine-paired lever was significantly reduced on sessions 6 and 7 relative to session 1 (Holm-Sidak, p<0.002, Cohen’s D>1.417).

We then tested the hypothesis that eliminating Sxc function would make rats more vulnerable to relapse. To do this, rats were injected with a sub-threshold dose of cocaine prior to a reinstatement test. This dose (3.0 mg/kg, IP) does not reinstate drug seeking in WT rats following extinction (figure 6E). Analyses of nonreinforced operant responding exhibited by WT and MSxc rats on the last extinction session and reinstatement test day revealed a significant interaction (F_1,28_=8.179, p=0.008, η_p_^2^ = 0.226). WT and MSxc did not differ during the last extinction session (T_28_=0.242, p=0.811), but MSxc rats exhibited significantly more lever presses during reinstatement testing than WT rats (T_28_=2.723, p=0.011, Cohen’s D=1.008). Furthermore, only MSxc rats reinstated previously extinguished operant responding as this behavior was significantly higher during the reinstatement test compared with their last extinction session (T_13_=-5.805, p<0.001, Cohen’s D=2.165). As expected, WT rats did not reinstate cocaine seeking (T_15_=-1.640, p=0.122). Therefore, these data demonstrate that MSxc rats have increased vulnerability to relapse to cocaine seeking.

## Discussion

The near-universal reliance on glutamate signaling for brain function illustrates the immense therapeutic potential and risk of targeting this system to treat CNS disorders. The difficulty of maintaining efficacy while limiting serious adverse effects is a key barrier in the rationale development of glutamatergic therapeutics. A question crucial to the future development of glutamatergic therapeutics is whether the glutamate signaling network includes functionally specialized components that regulate a limited, narrow range of brain function. Here we show using a novel genetically modified rat that a loss of astrocytic glutamate release via Sxc results in significant increases in approach behavior directed at a food-predictive cue in Pavlovian Conditioned Approach task and cocaine-primed drug seeking in the reinstatement paradigm. Both behaviors are used to model poor behavioral control in humans displaying impulsive or compulsive behaviors such as substance abuse (Garavan and Hester, 2007; Flagel et al., 2008; Connolly et al., 2012; Robinson et al., 2014). In contrast to neuronal glutamatergic mechanisms, a loss of Sxc was not fatal and did not produce impairments in development, physiological telemetry, basic learning, memory, or other indicators of widespread disruption of brain function. Hence, we propose that Sxc belongs to a class of unique, evolutionary new molecular adaptations that contributed to phylogenetic gains in behavior control.

Insights into Sxc function can be obtained from the evolutionary history of *Slc7a11/*xCT. The evolutionary origins of the key Sxc-subunit xCT occurred after the divergence of protostome and deuterostomes (Lewerenz et al., 2013). Here, we learned that *Slc7a11*/xCT is universally expressed in vertebrates, the majority of *Slc7a11*/xCT mutations did not alter the amino acid sequence of xCT protein indicating that altered protein sequence is selected against because of the potential for reduced biological fitness of the organism, the *Slc7a11*/xCT gene tree broadly matches the phylogenetic relationships among sampled vertebrate species (Irisarri et al., 2017), and the continued genetic divergence of *Slc7a11*/xCT may be further advancing the complexity of the mammalian brain. In support of the latter, we found potential Sxc regulatory sites, such as xCT_S26_, that were more likely to be present in primates than in birds, reptiles, fish, and in one instance, other mammals. Collectively, these results are similar to what has been found with other glutamatergic mechanisms that are known to be critical to brain function (Ryan and Grant, 2009; Teng et al., 2010; Moroz et al., 2014), which suggests that Sxc is a key glutamatergic signaling mechanism.

Next, we directly assessed the range of brain functions impacted by Sxc by creating a novel genetically modified rat lacking Sxc (MSxc). MSxc rats did not differ from WT rats on basic health and telemetry metrics; there were no genotypic differences in survival rates for up to one year, body weights between weaning and adulthood, core body temperatures, or general activity states. A loss of Sxc activity also did not impact basic learning or memory since WT and MSxc rats were equally adept in tasks involving classical conditioning, operant conditioning, extinction learning, object recognition memory, and discrimination of reward-paired stimuli. These findings are consistent with prior observations that xCT^-/-^ mice display only subtle changes in brain function (De Bundel et al., 2011; McCullagh and Featherstone, 2014; Williams and Featherstone, 2014), acute pharmacological inhibition of Sxc in rat impacts complex brain function such as cognitive set shifting without altering simple behaviors such as locomotor activity (Lutgen et al., 2014), and that chronic treatment with the Sxc enhancer N-acetylcysteine in humans is devoid of traditional CNS side effects (Pendyala and Creaven, 1995; Tenorio et al., 2021). Hence, a loss of Sxc activity does not produce the type of widespread changes in brain function observed with a loss of synaptic glutamate mechanisms (Tanaka et al., 1997; Wojcik et al., 2004; Christie et al., 2010; Kasthuri et al., 2015).

Further behavioral testing revealed that MSxc rats displayed higher levels of nonreinforced or negative-outcome behaviors despite the lack of widespread impairments in brain function. In Pavlovian Conditioned Approach, MSxc and WT rats appropriately increase their goal tray entries during the presentation of the CS+ but not the CS-. This demonstrates that WT and MSxc rats have an equivalent capacity to discriminate reward-paired stimuli. However, MSxc rats approached the CS+ more frequently than WT rats. Sensitized CS+ approach behavior in MSxc rats persisted even when the behavior was punished, which clearly indicates the maladaptive nature of the response. In the cocaine self-administration/reinstatement paradigm, there were no genotypic differences in acquisition, maintenance, or extinction of cocaine seeking behavior. However, MSxc rats displayed significantly higher levels of cocaine-primed drug seeking than WT rats. Heightened levels of sign-tracking and cocaine-primed reinstatement of drug seeking observed in MSxc rats have been used to model impulsive/compulsive behaviors in humans (Garavan and Hester, 2007; Flagel et al., 2008; Connolly et al., 2012; Robinson et al., 2014), indicating that the behavioral deficits observed in MSxc rats may reflect impaired behavioral control. Consistent with this, the Sxc enhancer N-acetylcysteine improves impaired cognitive control in human CNS disorders including trichotillomania (Grant et al., 2009), pathological gambling (Grant et al., 2007), substance abuse (Froeliger et al., 2015; Schulte et al., 2018; Woodcock et al., 2021), and obsessive-compulsive disorder (Oliver et al., 2015).

Considering our present findings along with past results obtained from xCT^-/-^ mice and N-acetylcysteine treatment in humans, we propose that Sxc is among the unique molecular adaptations with specialized functions that enabled phylogenetic gains in behavioral regulation. In part, we propose that the narrow range of brain functions requiring Sxc-mediated glutamate release reflects its relatively recent origin, which is hundreds of millions of years younger than synaptic mechanisms. As a result of its recent evolutionary origins, xCT is not expressed in most animals. In contrast, synaptic glutamate transmission is a highly ancient signaling mechanism that predated the emergence of complex nervous systems and likely exists in every animal (Ryan and Grant, 2009; Moroz et al., 2014; Moroz et al., 2021). Given that much of the brain evolved with synaptic glutamate in place, it is not surprising that it is a vital signaling process.

The possibility that evolutionary insights may be related to the range of functions reliant on a mechanism is supported by data obtained with GluN2 subunits of the NMDA receptor and the Dlg family of synaptic proteins (Emes et al., 2008; Nithianantharajah et al., 2013; Grant, 2016). Evolutionarily new paralogues of the Dlg family of synaptic proteins have been found to be selectively required for complex forms of behavior or cognition but not simpler forms of brain function (Nithianantharajah et al., 2013). Similarly, specialized cognitive abilities are functionally linked to evolutionarily newer adaptations to NMDA receptors (Ryan et al., 2008; Ryan et al., 2013). Collectively, these results indicate that evolutionarily new signaling mechanisms can have specialized roles in regulating behavior, which may expedite the development of novel approaches to treat behavioral control disorders.

A question that may be critical in better understanding the functional role of astrocytes is whether altered Sxc activity is impacting top-down behavioral control and/or bottom-up drivers of behavior. Ablation of the prefrontal cortex in primates and other animals mimics the behavioral pattern observed here in that goal tracking is replaced by sign tracking (Stepien, 1974), which illustrates that this pattern of behavior could reflect impaired cognitive control over behavior. If future work establishes that the deficits in MSxc rats reflects impaired cognitive control, then these data would add to the evidence indicating that astrocyte evolution contributed to the signaling complexity needed for highly sophisticated forms of cognition. Evidence consistent with this possibility include observations that individual astrocytes in the human cortex contact millions of synapses, considerably more than the estimated 30,000 synapses of cortical neurons (DeFelipe et al., 2002; Oberheim et al., 2006; Oberheim et al., 2009). Further, astrocytic glutamate impacts neuronal networks by coordinating communication between discrete neuronal ensembles (Lee et al., 2014; Martin et al., 2015; Poskanzer and Yuste, 2016; Sardinha et al., 2017). Lastly, enhanced complexity of spatial navigation information during seasonal migration are paralleled by increases in the number and morphological complexity of astrocytes within hippocampal networks (da Costa et al., 2020; Henrique et al., 2021; de Almeida Miranda et al., 2022). However, enhanced sign-tracking and cocaine reinstatement can arise from either increased bottom-up stimulation of behavior from processes such as incentive salience or reduced top-down regulation of behavior involving the frontal cortex and processes such as cognitive control (Jentsch and Taylor, 1999; Flagel et al., 2008; Feil et al., 2010; Robinson et al., 2014; Bickel et al., 2016; Kuhn et al., 2018; Sarter and Phillips, 2018; Gillis and Morrison, 2019; Colaizzi et al., 2020). Hence, determining the functional role of astrocytic glutamate signaling mechanisms can provide important insights into the organization structure of signaling within the vertebrate brain.

## Acknowledgements

This work was supported by DA035088 and DA050180, and grants from the JJ Keller Foundation and the Charles E. Kubly Mental Research Center.

## References

Back SE, McCauley JL, Korte KJ, Gros DF, Leavitt V, Gray KM, Hamner MB, DeSantis SM, Malcolm R, Brady KT, Kalivas PW (2016) A Double-Blind, Randomized, Controlled Pilot Trial of N-Acetylcysteine in Veterans With Posttraumatic Stress Disorder and Substance Use Disorders. J Clin Psychiatry 77:e1439–e1446.

Benson DA, Karsch-Mizrachi I, Lipman DJ, Ostell J, Sayers EW (2009) GenBank. Nucleic Acids Res 37:D26–31.

Bickel WK, Snider SE, Quisenberrys AJ, Stein JS, Hanlon CA (2016) Competing neurobehavioral decision systems theory of cocaine addiction: From mechanisms to therapeutic opportunities. Prog Brain Res 223:269–293.

Christie LA, Russell TA, Xu J, Wood L, Shepherd GM, Contractor A (2010) AMPA receptor desensitization mutation results in severe developmental phenotypes and early postnatal lethality. Proc Natl Acad Sci U S A 107:9412–9417.

Colaizzi JM, Flagel SB, Joyner MA, Gearhardt AN, Stewart JL, Paulus MP (2020) Mapping sign-tracking and goal-tracking onto human behaviors. Neurosci Biobehav Rev 111:84–94.

Connolly CG, Foxe JJ, Nierenberg J, Shpaner M, Garavan H (2012) The neurobiology of cognitive control in successful cocaine abstinence. Drug Alcohol Depend 121:45–53.

Cullen KR, Klimes-Dougan B, Westlund Schreiner M, Carstedt P, Marka N, Nelson K, Miller MJ, Reigstad K, Westervelt A, Gunlicks-Stoessel M, Eberly LE (2018) N-Acetylcysteine for Nonsuicidal Self-Injurious Behavior in Adolescents: An Open-Label Pilot Study. J Child Adolesc Psychopharmacol 28:136–144.

da Costa ER, Henrique EP, da Silva JB, Pereira PDC, de Abreu CC, Fernandes TN, Magalhães NGM, de Jesus Falcão da Silva A, Guerreiro LCF, Diniz CG, Diniz CWP, Diniz DG (2020) Changes in hippocampal astrocyte morphology of Ruddy turnstone (Arenaria interpres) during the wintering period at the mangroves of Amazon River estuary. J Chem Neuroanat 108:101805.

de Almeida Miranda D, Araripe J, de Morais Magalhães NG, de Siqueira LS, de Abreu CC, Pereira PDC, Henrique EP, da Silva Chira PAC, de Melo MAD, do Rêgo PS, Diniz DG, Sherry DF, Diniz CWP, Guerreiro-Diniz C (2022) Shorebirds’ Longer Migratory Distances Are Associated With Larger ADCYAP1 Microsatellites and Greater Morphological Complexity of Hippocampal Astrocytes. Frontiers in Psychology 12.

De Bundel D, Schallier A, Loyens E, Fernando R, Miyashita H, Van Liefferinge J, Vermoesen K, Bannai S, Sato H, Michotte Y, Smolders I, Massie A (2011) Loss of system x(c)-does not induce oxidative stress but decreases extracellular glutamate in hippocampus and influences spatial working memory and limbic seizure susceptibility. J Neurosci 31:5792–5803.

DeFelipe J, Alonso-Nanclares L, Arellano JI (2002) Microstructure of the neocortex: comparative aspects. J Neurocytol 31:299–316.

Delport W, Poon AF, Frost SD, Kosakovsky Pond SL (2010) Datamonkey 2010: a suite of phylogenetic analysis tools for evolutionary biology. Bioinformatics 26:2455–2457.

Durkee CA, Araque A (2019) Diversity and Specificity of Astrocyte-neuron Communication. Neuroscience 396:73–78.

Edgar RC (2004) MUSCLE: multiple sequence alignment with high accuracy and high throughput. Nucleic Acids Res 32:1792–1797.

Emes RD, Pocklington AJ, Anderson CN, Bayes A, Collins MO, Vickers CA, Croning MD, Malik BR, Choudhary JS, Armstrong JD, Grant SG (2008) Evolutionary expansion and anatomical specialization of synapse proteome complexity. Nat Neurosci 11:799–806.

Fairless R, Bading H, Diem R (2021) Pathophysiological Ionotropic Glutamate Signalling in Neuroinflammatory Disease as a Therapeutic Target. Front Neurosci 15:741280.

Feil J, Sheppard D, Fitzgerald PB, Yucel M, Lubman DI, Bradshaw JL (2010) Addiction, compulsive drug seeking, and the role of frontostriatal mechanisms in regulating inhibitory control. Neurosci Biobehav Rev 35:248–275.

Flagel SB, Watson SJ, Akil H, Robinson TE (2008) Individual differences in the attribution of incentive salience to a reward-related cue: influence on cocaine sensitization. Behav Brain Res 186:48–56.

Froeliger B, McConnell PA, Stankeviciute N, McClure EA, Kalivas PW, Gray KM (2015) The effects of N-Acetylcysteine on frontostriatal resting-state functional connectivity, withdrawal symptoms and smoking abstinence: A double-blind, placebo-controlled fMRI pilot study. Drug Alcohol Depend 156:234–242.

Garavan H, Hester R (2007) The role of cognitive control in cocaine dependence. Neuropsychol Rev 17:337–345.

Geurts AM, Cost GJ, Remy S, Cui X, Tesson L, Usal C, Menoret S, Jacob HJ, Anegon I, Buelow R (2010) Generation of gene-specific mutated rats using zinc-finger nucleases. Methods Mol Biol 597:211–225.

Gillis ZS, Morrison SE (2019) Sign Tracking and Goal Tracking Are Characterized by Distinct Patterns of Nucleus Accumbens Activity. eNeuro 6.

Grant JE, Kim SW, Odlaug BL (2007) N-acetyl cysteine, a glutamate-modulating agent, in the treatment of pathological gambling: a pilot study. Biol Psychiatry 62:652–657.

Grant JE, Odlaug BL, Kim SW (2009) N-acetylcysteine, a glutamate modulator, in the treatment of trichotillomania: a double-blind, placebo-controlled study. Archives of general psychiatry 66:756–763.

Grant JE, Chamberlain SR, Redden SA, Leppink EW, Odlaug BL, Kim SW (2016) N-Acetylcysteine in the Treatment of Excoriation Disorder: A Randomized Clinical Trial. JAMA Psychiatry 73:490–496.

Grant SG (2016) The molecular evolution of the vertebrate behavioural repertoire. Philos Trans R Soc Lond B Biol Sci 371:20150051.

Harrison RA, Wefel JS (2018) Neurocognitive Function in Adult Cancer Patients. Neurol Clin 36:653–674.

Henrique EP, de Oliveira MA, Paulo DC, Pereira PDC, Dias C, de Siqueira LS, de Lima CM, Miranda DA, do Rego PS, Araripe J, de Melo MAD, Diniz DG, de Morais Magalhães NG, Sherry DF, Picanço Diniz CW, Diniz CG (2021) Contrasting migratory journeys and changes in hippocampal astrocyte morphology in shorebirds. Eur J Neurosci 54:5687–5704.

Henter ID, Park LT, Zarate CA, Jr. (2021) Novel Glutamatergic Modulators for the Treatment of Mood Disorders: Current Status. CNS Drugs 35:527–543.

Horner AE, Heath CJ, Hvoslef-Eide M, Kent BA, Kim CH, Nilsson SR, Alsio J, Oomen CA, Holmes A, Saksida LM, Bussey TJ (2013) The touchscreen operant platform for testing learning and memory in rats and mice. Nature protocols 8:1961–1984.

Hyman SE, Malenka RC, Nestler EJ (2006) Neural mechanisms of addiction: the role of reward-related learning and memory. Annu Rev Neurosci 29:565–598.

Irisarri I, Baurain D, Brinkmann H, Delsuc F, Sire JY, Kupfer A, Petersen J, Jarek M, Meyer A, Vences M, Philippe H (2017) Phylotranscriptomic consolidation of the jawed vertebrate timetree. Nat Ecol Evol 1:1370–1378.

Jentsch JD, Taylor JR (1999) Impulsivity resulting from frontostriatal dysfunction in drug abuse: implications for the control of behavior by reward-related stimuli. Psychopharmacology (Berl) 146:373–390.

Kasthuri N et al. (2015) Saturated Reconstruction of a Volume of Neocortex. Cell 162:648–661.

Kosakovsky Pond SL, Frost SD (2005) Not so different after all: a comparison of methods for detecting amino acid sites under selection. Mol Biol Evol 22:1208–1222.

Koshy Cherian A, Kucinski A, Pitchers K, Yegla B, Parikh V, Kim Y, Valuskova P, Gurnani S, Lindsley CW, Blakely RD, Sarter M (2017) Unresponsive Choline Transporter as a Trait Neuromarker and a Causal Mediator of Bottom-Up Attentional Biases. J Neurosci 37:2947–2959.

Kuhn BN, Campus P, Flagel SB, Tomie A, Morrow J (2018) The neurobiological mechanisms underlying sign-tracking behavior. Sign-tracking and drug addiction.

Lange F, Seer C, Kopp B (2017) Cognitive flexibility in neurological disorders: Cognitive components and event-related potentials. Neurosci Biobehav Rev 83:496–507.

Lee HS, Ghetti A, Pinto-Duarte A, Wang X, Dziewczapolski G, Galimi F, Huitron-Resendiz S, Pina-Crespo JC, Roberts AJ, Verma IM, Sejnowski TJ, Heinemann SF (2014) Astrocytes contribute to gamma oscillations and recognition memory. Proc Natl Acad Sci U S A 111:E3343–3352.

Lewerenz J, Hewett SJ, Huang Y, Lambros M, Gout PW, Kalivas PW, Massie A, Smolders I, Methner A, Pergande M, Smith SB, Ganapathy V, Maher P (2013) The cystine/glutamate antiporter system x(c)(-) in health and disease: from molecular mechanisms to novel therapeutic opportunities. Antioxid Redox Signal 18:522–555.

Lovic V, Saunders BT, Yager LM, Robinson TE (2011) Rats prone to attribute incentive salience to reward cues are also prone to impulsive action. Behav Brain Res 223:255–261.

Luscher C, Robbins TW, Everitt BJ (2020) The transition to compulsion in addiction. Nat Rev Neurosci 21:247–263.

Lutgen V, Resch J, Qualmann K, Raddatz NJ, Panhans C, Olander EM, Kong L, Choi S, Mantsch JR, Baker DA (2014) Behavioral assessment of acute inhibition of system xc (-) in rats. Psychopharmacology (Berl) 231:4637–4647.

Madayag A, Kau KS, Lobner D, Mantsch JR, Wisniewski S, Baker DA (2010) Drug-induced plasticity contributing to heightened relapse susceptibility: neurochemical changes and augmented reinstatement in high-intake rats. J Neurosci 30:210–217.

Martin R, Bajo-Graneras R, Moratalla R, Perea G, Araque A (2015) Circuit-specific signaling in astrocyte-neuron networks in basal ganglia pathways. Science 349:730–734.

McCullagh EA, Featherstone DE (2014) Behavioral characterization of system xc-mutant mice. Behav Brain Res 265:1–11.

McCutcheon JE, Marinelli M (2009) Age matters. Eur J Neurosci 29:997–1014.

Moore BR (2004) The evolution of learning. Biol Rev Camb Philos Soc 79:301–335.

Moroz LL, Nikitin MA, Policar PG, Kohn AB, Romanova DY (2021) Evolution of glutamatergic signaling and synapses. Neuropharmacology 199:108740.

Moroz LL et al. (2014) The ctenophore genome and the evolutionary origins of neural systems. Nature 510:109–114.

Nithianantharajah J, Komiyama NH, McKechanie A, Johnstone M, Blackwood DH, St Clair D, Emes RD, van de Lagemaat LN, Saksida LM, Bussey TJ, Grant SG (2013) Synaptic scaffold evolution generated components of vertebrate cognitive complexity. Nat Neurosci 16:16–24.

Oberheim NA, Wang X, Goldman S, Nedergaard M (2006) Astrocytic complexity distinguishes the human brain. Trends Neurosci 29:547–553.

Oberheim NA, Takano T, Han X, He W, Lin JH, Wang F, Xu Q, Wyatt JD, Pilcher W, Ojemann JG, Ransom BR, Goldman SA, Nedergaard M (2009) Uniquely hominid features of adult human astrocytes. J Neurosci 29:3276–3287.

Oliver G, Dean O, Camfield D, Blair-West S, Ng C, Berk M, Sarris J (2015) N-acetyl cysteine in the treatment of obsessive compulsive and related disorders: a systematic review. Clin Psychopharmacol Neurosci 13:12–24.

Ottestad-Hansen S, Hu QX, Follin-Arbelet VV, Bentea E, Sato H, Massie A, Zhou Y, Danbolt NC (2018) The cystine-glutamate exchanger (xCT, Slc7a11) is expressed in significant concentrations in a subpopulation of astrocytes in the mouse brain. Glia.

Paydary K, Akamaloo A, Ahmadipour A, Pishgar F, Emamzadehfard S, Akhondzadeh S (2016) N-acetylcysteine augmentation therapy for moderate-to-severe obsessive-compulsive disorder: randomized, double-blind, placebo-controlled trial. J Clin Pharm Ther 41:214–219.

Pendyala L, Creaven PJ (1995) Pharmacokinetic and pharmacodynamic studies of N-acetylcysteine, a potential chemopreventive agent during a phase I trial. Cancer Epidemiol Biomarkers Prev 4:245–251.

Peters R, Ee N, Peters J, Beckett N, Booth A, Rockwood K, Anstey KJ (2019) Common risk factors for major noncommunicable disease, a systematic overview of reviews and commentary: the implied potential for targeted risk reduction. Ther Adv Chronic Dis 10:2040622319880392.

Poskanzer KE, Yuste R (2016) Astrocytes regulate cortical state switching in vivo. Proc Natl Acad Sci U S A 113:E2675–2684.

Ramos-Vicente D, Ji J, Gratacos-Batlle E, Gou G, Reig-Viader R, Luis J, Burguera D, Navas-Perez E, Garcia-Fernandez J, Fuentes-Prior P, Escriva H, Roher N, Soto D, Bayes A (2018) Metazoan evolution of glutamate receptors reveals unreported phylogenetic groups and divergent lineage-specific events. Elife 7.

Robinson TE, Yager LM, Cogan ES, Saunders BT (2014) On the motivational properties of reward cues: Individual differences. Neuropharmacology 76 Pt B:450–459.

Ryan TJ, Grant SG (2009) The origin and evolution of synapses. Nat Rev Neurosci 10:701–712.

Ryan TJ, Emes RD, Grant SG, Komiyama NH (2008) Evolution of NMDA receptor cytoplasmic interaction domains: implications for organisation of synaptic signalling complexes. BMC Neurosci 9:6.

Ryan TJ, Kopanitsa MV, Indersmitten T, Nithianantharajah J, Afinowi NO, Pettit C, Stanford LE, Sprengel R, Saksida LM, Bussey TJ, O’Dell TJ, Grant SG, Komiyama NH (2013) Evolution of GluN2A/B cytoplasmic domains diversified vertebrate synaptic plasticity and behavior. Nat Neurosci 16:25–32.

Sardinha VM, Guerra-Gomes S, Caetano I, Tavares G, Martins M, Reis JS, Correia JS, Teixeira-Castro A, Pinto L, Sousa N, Oliveira JF (2017) Astrocytic signaling supports hippocampal-prefrontal theta synchronization and cognitive function. Glia 65:1944–1960.

Sarter M, Phillips KB (2018) The neuroscience of cognitive-motivational styles: Sign-and goal-trackers as animal models. Behav Neurosci 132:1–12.

Schulte MHJ, Wiers RW, Boendermaker WJ, Goudriaan AE, van den Brink W, van Deursen DS, Friese M, Brede E, Waters AJ (2018) The effect of N-acetylcysteine and working memory training on cocaine use, craving and inhibition in regular cocaine users: correspondence of lab assessments and Ecological Momentary Assessment. Addict Behav 79:24–31.

Squeglia LM, Tomko RL, Baker NL, McClure EA, Book GA, Gray KM (2018) The effect of N-acetylcysteine on alcohol use during a cannabis cessation trial. Drug Alcohol Depend 185:17–22.

Stepien I (1974) The magnet reaction, a symptom of prefrontal ablation. Acta Neurobiol Exp (Wars) 34:145–160.

Tamura K, Stecher G, Peterson D, Filipski A, Kumar S (2013) MEGA6: Molecular Evolutionary Genetics Analysis version 6.0. Mol Biol Evol 30:2725–2729.

Tanaka K, Watase K, Manabe T, Yamada K, Watanabe M, Takahashi K, Iwama H, Nishikawa T, Ichihara N, Kikuchi T, Okuyama S, Kawashima N, Hori S, Takimoto M, Wada K (1997) Epilepsy and exacerbation of brain injury in mice lacking the glutamate transporter GLT-1. Science 276:1699–1702.

Teng H, Cai W, Zhou L, Zhang J, Liu Q, Wang Y, Dai W, Zhao M, Sun Z (2010) Evolutionary mode and functional divergence of vertebrate NMDA receptor subunit 2 genes. PLoS One 5:e13342.

Tenorio M, Graciliano NG, Moura FA, Oliveira ACM, Goulart MOF (2021) N-Acetylcysteine (NAC): Impacts on Human Health. Antioxidants (Basel) 10.

Tunstall BJ, Kearns DN (2015) Sign-tracking predicts increased choice of cocaine over food in rats. Behavioural brain research 281:222–228.

Vandame D, Ulmann L, Teigell M, Prieto-Cappellini M, Vignon J, Privat A, Perez-Polo R, Nesic O, Hirbec H (2013) Development of NMDAR antagonists with reduced neurotoxic side effects: a study on GK11. PLoS One 8:e81004.

Verdejo-Garcia A, Clark L, Verdejo-Roman J, Albein-Urios N, Martinez-Gonzalez JM, Gutierrez B, Soriano-Mas C (2015) Neural substrates of cognitive flexibility in cocaine and gambling addictions. Br J Psychiatry 207:158–164.

Versaggi CL, King CP, Meyer PJ (2016) The tendency to sign-track predicts cue-induced reinstatement during nicotine self-administration, and is enhanced by nicotine but not ethanol. Psychopharmacology 233:2985–2997.

Volkow ND, Wise RA, Baler R (2017) The dopamine motive system: implications for drug and food addiction. Nat Rev Neurosci 18:741–752.

Williams LE, Featherstone DE (2014) Regulation of hippocampal synaptic strength by glial xCT. J Neurosci 34:16093–16102.

Wojcik SM, Rhee JS, Herzog E, Sigler A, Jahn R, Takamori S, Brose N, Rosenmund C (2004) An essential role for vesicular glutamate transporter 1 (VGLUT1) in postnatal development and control of quantal size. Proc Natl Acad Sci U S A 101:7158–7163.

Woodcock EA, Lundahl LH, Khatib D, Stanley JA, Greenwald MK (2021) N-acetylcysteine reduces cocaine-seeking behavior and anterior cingulate glutamate/glutamine levels among cocaine-dependent individuals. Addict Biol 26:e12900.

